# Known phyla dominate the RNA virome in Tara Oceans data, not new phyla as claimed

**DOI:** 10.1101/2022.07.30.502162

**Authors:** Robert C. Edgar

## Abstract

The Tara Oceans project claimed to identify five new RNA virus phyla, two of which “are dominant in the oceans”. However, their own assignments classify 28,353 of their putative RdRp-containing contigs to known phyla but only 886 (2.8%) to the five claimed new phyla combined. I mapped their reads to their contigs, finding that known phyla also account for a large majority (93.8%) of reads. I show that 510 of their putative viral contigs contain cellular proteins and further that predicted polymerase structures for their claimed phyla have incomplete and malformed palm domains, contradicting viral polymerase function and calling into question whether their putative polymerase sequences are derived from viruses.

## Introduction

In 2017, fewer than 5,000 RNA virus species were known (Wolf et al., 2018). Since then, high-throughput metagenomics has dramatically expanded the known RNA virome with studies such as Yangshan Deep Water Harbour (Wolf et al., 2020), Serratus (Edgar et al., 2022) and Tara Oceans (Zayed et al., 2022) collectively adding more than 140,000 new species. These new viruses are classified primarily on the basis of their RNA-dependent RNA polymerase (RdRp) gene sequences, with clusters at 90% sequence identity (viral Operational Taxonomic Units, vOTUs) serving as proxies for species.

### Tara Oceans data and analysis

The Tara Oceans raw data comprises metagenomic sequencing reads for 1,703 RNA-seq runs (Bio-Project PRJEB402). Reads were assembled, and 44,799 RdRp-containing (RdRp+) contigs were identified by hidden Markov model (HMM) search. Contigs were clustered at 90% nucleotide (nt) identity, yielding 5,504 vOTUs. Reads were mapped to RdRp+ contigs using bowtie2 (Langmead and Salzberg, 2012). Contig abundances per sample were calculated from the bowtie2 mapping using Cov-erM (https://github.com/wwood/CoverM).

### Claims made by the Tara study

Tara claimed to identify five novel RNA virus phyla from sequence analysis of this data, with no corroborating evidence from traditional methods such as isolated or cultured viruses or electron microscopy of virions. According to their Abstract, “‘[s]pecies’-rank abundance determination revealed that viruses of the new phyla ‘Taraviricota’ … and ‘Arctiviricota’ are widespread and dominant in the oceans”. See my Fig. 1 for this claim in context.

**Figure 1.**
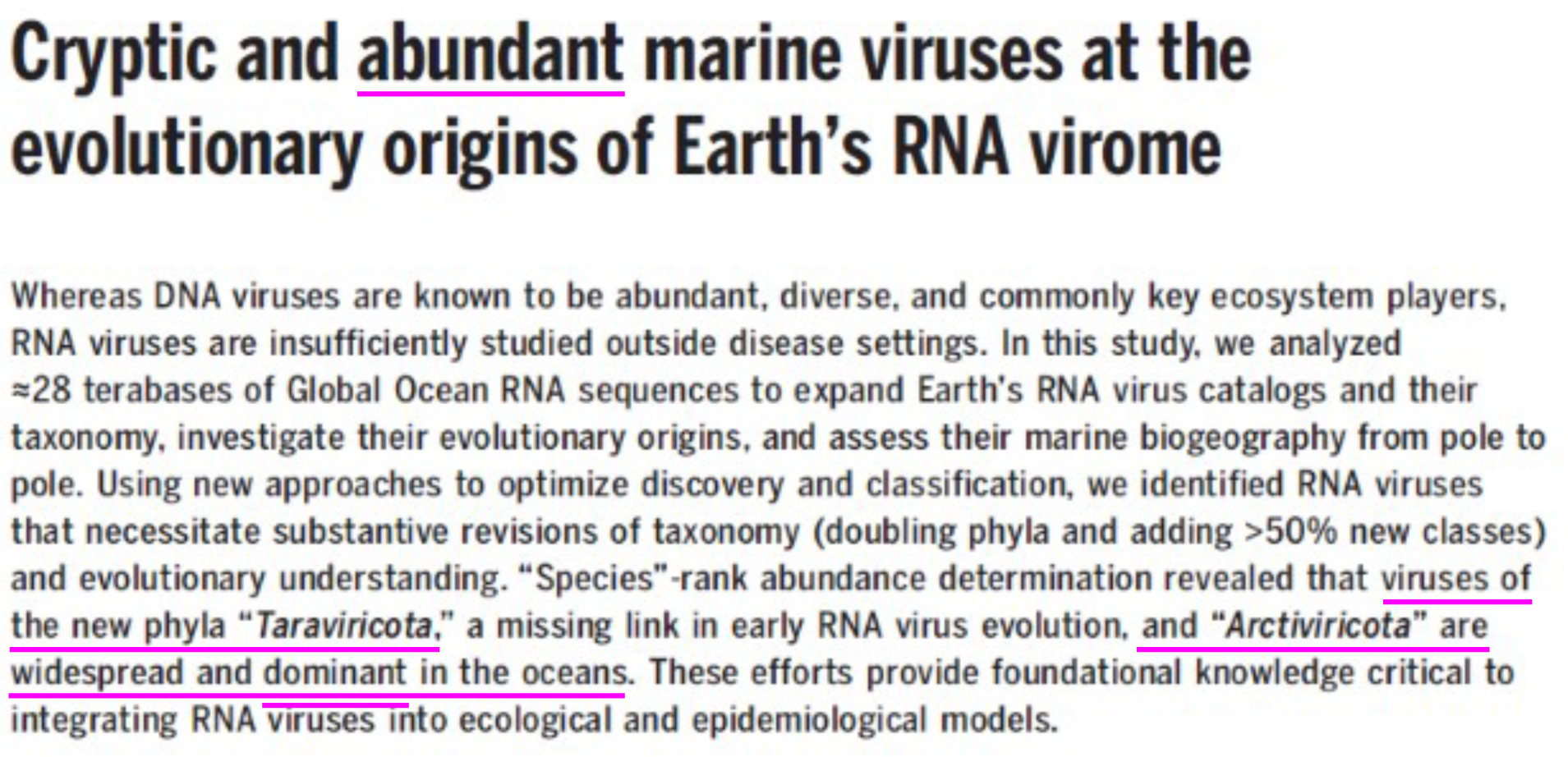
Main claim of the Tara paper. Magenta colored underlines added for emphasis. The title describes “cryptic” (i.e., not well-characterized) marine viruses as “abundant”, which implies a notably high abundance given its appearance in a short title. The Abstract claims more specifically that “Taraviricota” and “Arctiviricota” are “dominant in the oceans”.

### Missing supporting data

Given the claim that novel viruses dominate the oceans by abundance, I would expect to see a table giving vOTU abundances per sample, but no such table is provided. Further, sufficient data and code required to independently check their abundance calculations are not provided in the supplementary materials or associated data repositories. Assemblies are deposited, but metadata associating a contig with its sequencing run is provided only for a small subset (35%) of contigs. Abundances per sample per contig as reported by CoverM are not provided, and the code required to calculate per-”megataxon” abundances from per-contig / per-sample abundances is also not provided.

To implement an independent assessment of vOTU abundances across the complete Tara dataset, I therefore had to start from scratch by mapping reads and writing my own analysis scripts. Code and intermediate data supporting my re-analysis are provided in my supplementary materials, Zenodo data repository and github.

### Known phyla dominate Tara’s contigs

The number of contigs per megataxon is shown in my Table 1, obtained by a one-line Linux command from their Table S6:

~~~
  grep “This study” TableS6.tsv | cut -f8 | sort | uniq -c | sort -nr
~~~

**Table 1.** Contig classifications per Tara’s Table S6. The numbers in this table were generated from Tara’s supplementary table S6 using a one-line Linux command given in my main text under ‘Known phyla dominate Tara’s contigs’. This shows that known phyla account for a large majority of contigs, while the five claimed novel phyla combined account for only 1.98%.

| *Mega-taxon* | *Mega-cluster* | *Rank* | *Contigs* | *Pct. contigs* |
| --- | --- | --- | --- | --- |
| (Not determined) | (None) | (None) | 13436 | 30.0% |
| Pisuviricota | 1 | Phylum | 13014 | 29.0% |
| Kitrinoviricota | 2 | Phylum | 5543 | 12.4% |
| Lenarviricota;others | 3 | Split phylum | 7060 | 15.8% |
| Lenarviricota;Allassoviricetes | 4 | Class | 154 | 0.34% |
| Negarnaviricota;Polyploviricotina | 5 | Sub-phylum | 1133 | 2.53% |
| Duplornaviricota;Chrymotiviricetes | 6 | Class | 655 | 1.46% |
| Negarnaviricota;Haploviricotina | 7 | Sub-phylum | 528 | 1.18% |
| Wei-like | 8 | ? | 1971 | 4.40% |
| Duplornaviricota;Spinareovirinae | 9 | Sub-family | 0 | 0% |
| Permuto-like | 10 | ? | 118 | 0.26% |
| Duplornaviricota;Sedoreovirinae | 11 | Sub-family | 237 | 0.53% |
| **New_12at1.1 (Taraviricota)** | **12** | **"Phylum"** | **644** | **1.44%** |
| Kitrino-like | 13 | ? | 40 | 0.09% |
| **New_14at1.1 (Pomiviricota)** | **14** | **"Phylum"** | **119** | **0.27%** |
| Duplornaviricota;Vidaverviricetes | 15 | Class | 29 | 0.06% |
| **New_16at1.1 (Paraxenoviricota)** | **16** | **"Phylum"** | **14** | **0.03%** |
| **New_17at1.1 (Lenar-like)** | 17 | ? | 24 | 0.05% |
| **New_18at1.1 (Wamoviricota)** | **18** | **"Phylum"** | **4** | **0.01%** |
| **New_19at1.1 (Arctiviricota)** | **19** | **"Phylum"** | **105** | **0.23%** |

This shows that “Taraviricota” and “Arctiviricota” account for 1.44% and 0.23% of their contigs, respectively, contradicting their claim that these viruses “dominate”. Known phyla together account for 84.1% of contigs (excluding “not determined”) and thereby dominate Tara’s RNA virome as measured by total numbers of contigs.

### Known phyla dominate Tara’s reads

To map reads for all Tara samples to vOTUs, I used Serratus to launch a cluster of cloud CPUs running the diamond (Buchfink et al., 2015) read mapper. “Megataxon” assignments of vOTUs were taken from Tara supplementary tables and used to assign each vOTU to a phylum. The total number of reads mapped to each phylum is shown in my Table 2, showing that “Taraviricota” and “Arctiviricota” account for 3.75% and 1.28% of the mapped reads, respectively. Read frequency per phylum correlates well with contig frequency per phylum (*r*=0.79) as would be expected. Otherwise, read depth per phylum would necessarily be highly skewed, demanding further investigation and discussion. In fact, known phyla together account for 93.8% of mapped reads (excluding contigs annotated as “not determined” by Tara), and thereby dominate Tara’s RNA virome as measured by total numbers of mapped reads.

**Table 2.** Numbers of mapped reads across the complete Tara RNA virome. This table shows the total numbers of reads mapped to each phylum by Serratus from all Tara RNA-seq runs. Phylum classifications were taken from Tara’s Tables S6 and S7. The number of reads mapped to each phylum correlates with the number of contigs (Table 1, *r* = 0.79) as would be expected. Thus, known phyla dominate Tara’s reads.

| Phylum | Nr. of mapped reads | Pct. of mapped reads |
| --- | --- | --- |
| *Pisuviricota* | 4,948,767 | 48.0% |
| *Kitrinaviricota* | 1,880,720 | 18.3% |
| *Lenarviricota* | 1,423,964 | 13.8% |
| (Undetermined) | 1,071,627 | 10.4% |
| "Taraviricota" | 385,966 | 3.75% |
| *Negarnaviricota* | 349,901 | 3.40% |
| "Arctiviricota" | 132,257 | 1.28% |
| *Duplornaviricota* | 49,879 | 0.48% |
| "Pomiviricota" | 37,470 | 0.36% |
| "Paraxenoviricota" | 17,417 | 0.17% |
| "Wamoviricota" | 789 | 0.01% |

### Known phyla dominate the Arctic Ocean

On p.6, third column of their paper, Tara claim that “vOTUs belonging to the -ssRNA phylum “Arctiviricota” were, on average, the most abundant across most of the Atlantic Arctic waters (Fig. 4).” My Fig. 2 shows a region of the Arctic Ocean where Tara’s Fig. 4 displays the highest abundances for “Arctiviricota”. In stark contrast, “Arctiviricota” accounts for only 0.01% of reads mapped by Serratus in that region.

**Figure 2.**
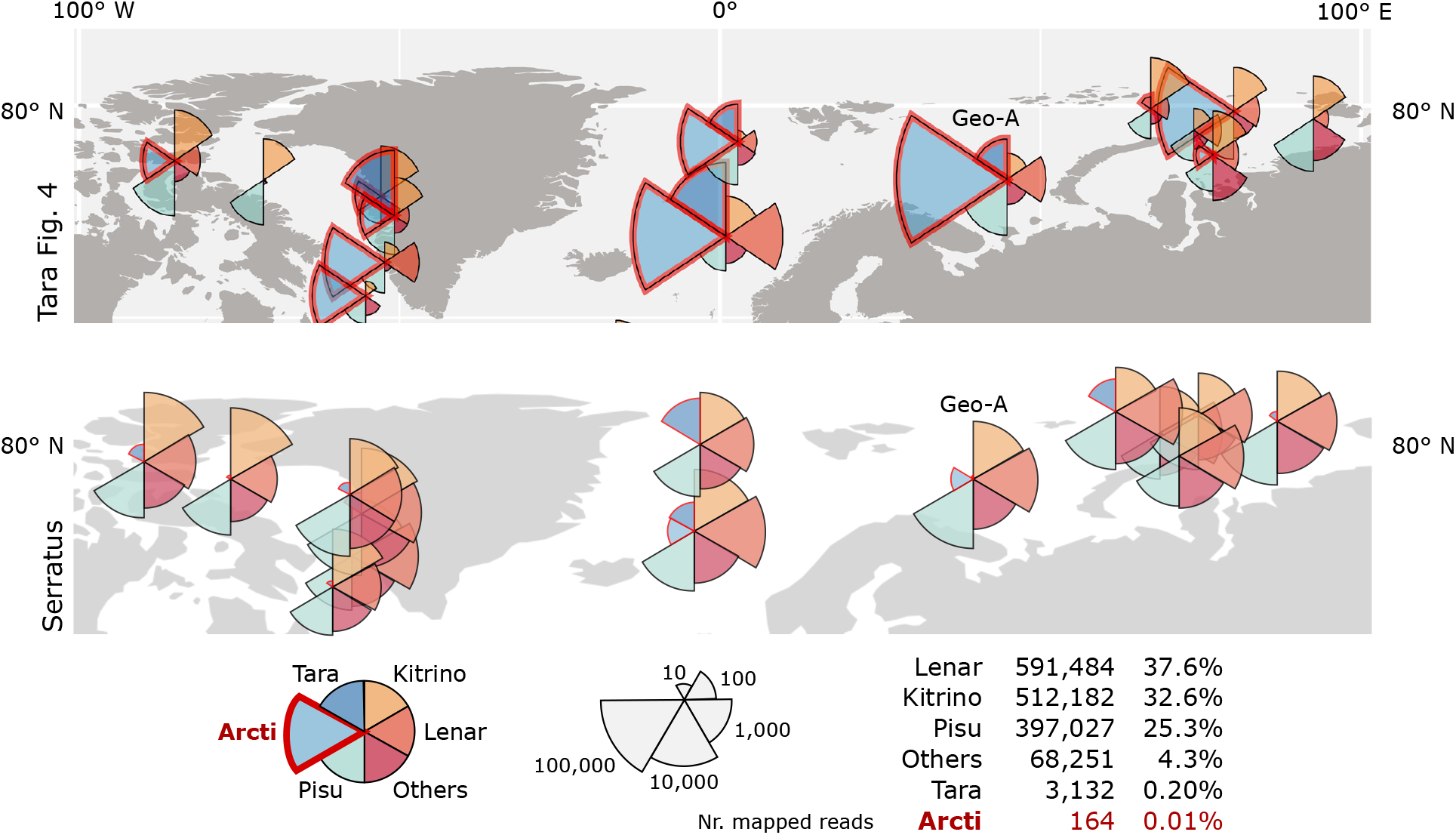
Phylum abundances in the Arctic Ocean. The figure shows abundances in the region bounded by 100°W to 100°E longitude and 60°N to 90°N latitude. Top is this region as depicted in Tara’s Fig. 4. Below is the same region with abundances measured as the numbers of reads mapped by Serratus. Reads are classified into six groups Tara (“Taraviricota”), Kitrino (*Kitrinoviricota*), etc., following Tara’s Fig. 4. My pie charts show the number of reads mapped to each group on a logarithmic scale such that the radius of the segment is proportional to log_10_*n* where *n* is the number of reads mapped to the group. I do not understand the scaling used in Tara’s pie charts and therefore did not attempt to reproduce them exactly. The total numbers of reads mapped in this region is shown at lower right, with only 164 reads assigned to “Arctiviricota” (0.01%). Location Geo-A shows high abundances of both “Taraviricota” and “Arctiviricota” in Tara’s Fig. 4; this location is analyzed in more detail in my Fig. 3 and my Table 4.

### Focused re-analysis of selected geolocations

My findings of low abundances for “Taraviricota” and “Arctiviricota” in contigs and reads contradict Tara’s main claim and raise the question of how the discrepancy might be explained. This question is difficult to address given the lack of supporting methods descriptions, data and code related to this claim. To the best of my knowledge, Tara’s Fig. 4 is the only place where they report abundances in any form, and I therefore selected four geolocations from this figure for further investigation: 72.5°N,44.1°E (Geo-A), 71.6°N,160.9°E (Geo-B), 71.1°N,175.0°E (Geo-C), and 21.1°S,104.8°W (Geo-D) (my Table 3). Geo-A was selected because it shows the largest abundance of “Arctiviricota” of all locations together with a large abundance of “Taraviricota”, and should therefore clearly show high abundances of both these “phyla” by any reasonable abundance metric. Geo-D was selected because it is shown as having one of the highest abundances of “Taraviricota”. Geo-B and Geo-C were selected as controls which have no visible abundance of “Taraviricota” and “Arctiviricota” in Tara’s Fig. 4.

**Table 3.** Selected geolocations. This table shows sample numbers, latitude (Lat.), longitude (Lon.) and SRA sequencing run accessions for the four geolocations selected for focused analysis. See also my Fig. 3.

| *Location* | *Sample* | *Lat.* | *Lon.* | *Runs* |
| --- | --- | --- | --- | --- |
| Geo-A | SAMEA4397209 | 72.5199 | 44.0552 | ERR3586697 ERR3586749 |
|  | SAMEA4397168 | 72.5432 | 44.1020 | ERR3586696 ERR3586718 |
|  | SAMEA4397112 | 72.5128 | 44.0775 | ERR3586695 ERR3586748 |
| Geo-B | SAMEA4397768 | 71.5955 | 160.9383 | ERR3586682 ERR3586742 |
| Geo-C | SAMEA4397809 | 71.0704 | 174.9916 | ERR3586683 ERR3586743 |
| Geo-D | SAMEA2622078 | -21.1460 | -104.7870 | ERR3586898 ERR3587004 |

I used four distinctly different read-mapping procedures to check that results are consistent and robust against anomalies in any single method (in particular, unusually large numbers of false-positive or false-negative alignments), as follows. (1) My own re-implementation of Tara’s read mapping with bowtie2 where the reference database is nt sequences of all RdRp+ contigs; both reads and contigs were poly-A-trimmed by bbduk (https://jgi.doe.gov/data-and-tools/bbtools/) before mapping. (2) Serratus with its diamond aligner module, as used in mapping of the full Tara dataset. (3) The diamond aligner using standard NCBI tools to download reads. Method (2) generates identical results to (1) but is more computationally expensive. Methods (2) and (3) used a vOTU reference comprising 5,546 amino acid sequences trimmed to approximately full-length RdRp domains and clustered at 90% identity. (4) The diamond aligner using a reference constructed by 6-frame translation of all RdRp+ contigs, to check how many additional reads are mapped by diamond when non-RdRp sequence is retained and clustering is not used to reduce redundancy in the reference. I used 6-frame translation to ensure that all valid CDS is included (as opposed to ORF-finding or other methods designed to find CDS and discard non-CDS) because this method is robust against frame-shifts and non-standard genetic codes up to a small fraction of mis-translated codons which should have minimal impact on measured coverage. The total data size of these four geolocations combined is 120Gb, enabling download and mapping on a single Linux server in a few hours and thereby facilitating independent verification of my results.

Abundances measured as number of reads mapped are summarized in my Fig. 3 and Table 4. My Fig. 3 shows that the five methods give a consistent qualitative picture of the abundances of the six groups used in Tara Fig. 4. My Table 4 shows that the largest fraction of reads mapped to “Arctiviricota” at Geo-A is 68/445,728 = 0.015% by Serratus, and no reads mapped to “Taraviricota” at any of the four geolocations by any of the five methods. This definitively contradicts the pictures shown at Geo-A and Geo-D in Tara Fig. 4, regardless of the chosen abundance measure.

**Figure 3.**
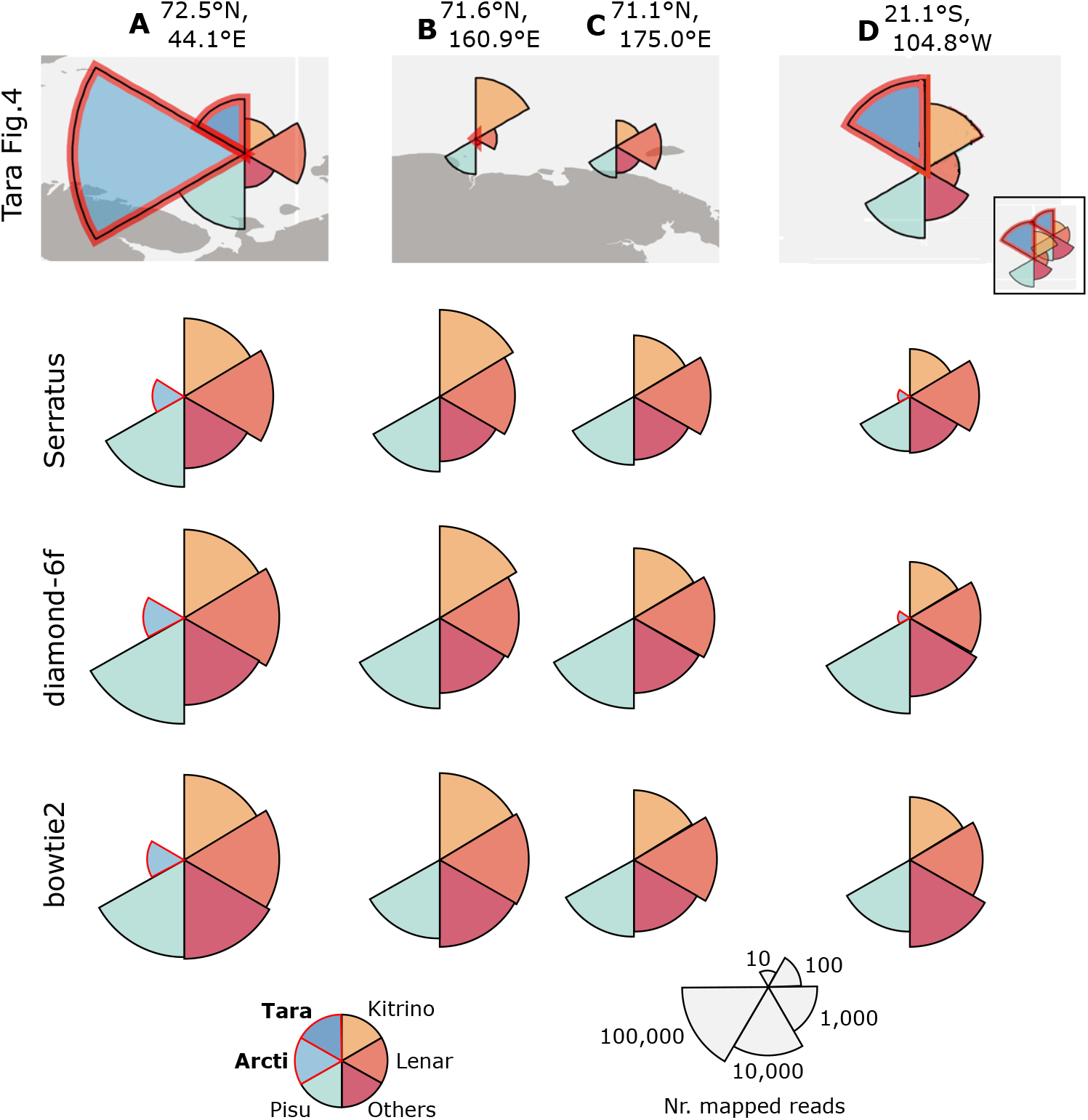
Phylum abundances at four selected geolocations. Abundances are shown as pie charts in a four-by-four grid. Columns are locations Geo-A, Geo-B, Geo-C and Geo-D as specified in my Table 3. The top row shows the locations as depicted in Tara’s Fig. 4. The pie-chart at location Geo-D was manually edited to remove the overlapping location adjacent to it (seen in its unedited form in the small inset). The three remaining rows show my own read mapping results using three different methods: Serratus is method (1), diamond-6f is method (3) and bowtie2 is method (4) as described in my main text under ‘Focused re-analysis of selected geolocations’. Reads are classified into six groups Tara (“Taraviricota”), Kitrino (*Kitrinoviricota*), etc., following Tara’s Fig. 4. My pie charts show the number of reads mapped to each group on a logarithmic scale such that the radius of the segment is proportional to *log_10_n* where *n* is the number of reads mapped to the group. I do not understand the scaling used in Tara’s pie charts and therefore did not attempt to reproduce them exactly. However, all my methods agree that exactly one read mapped to “Taraviricota” at Geo-A and none at Geo-D, and my results therefore contradict their figure regardless of their scaling. See also my Table 4.

**Figure 4.**
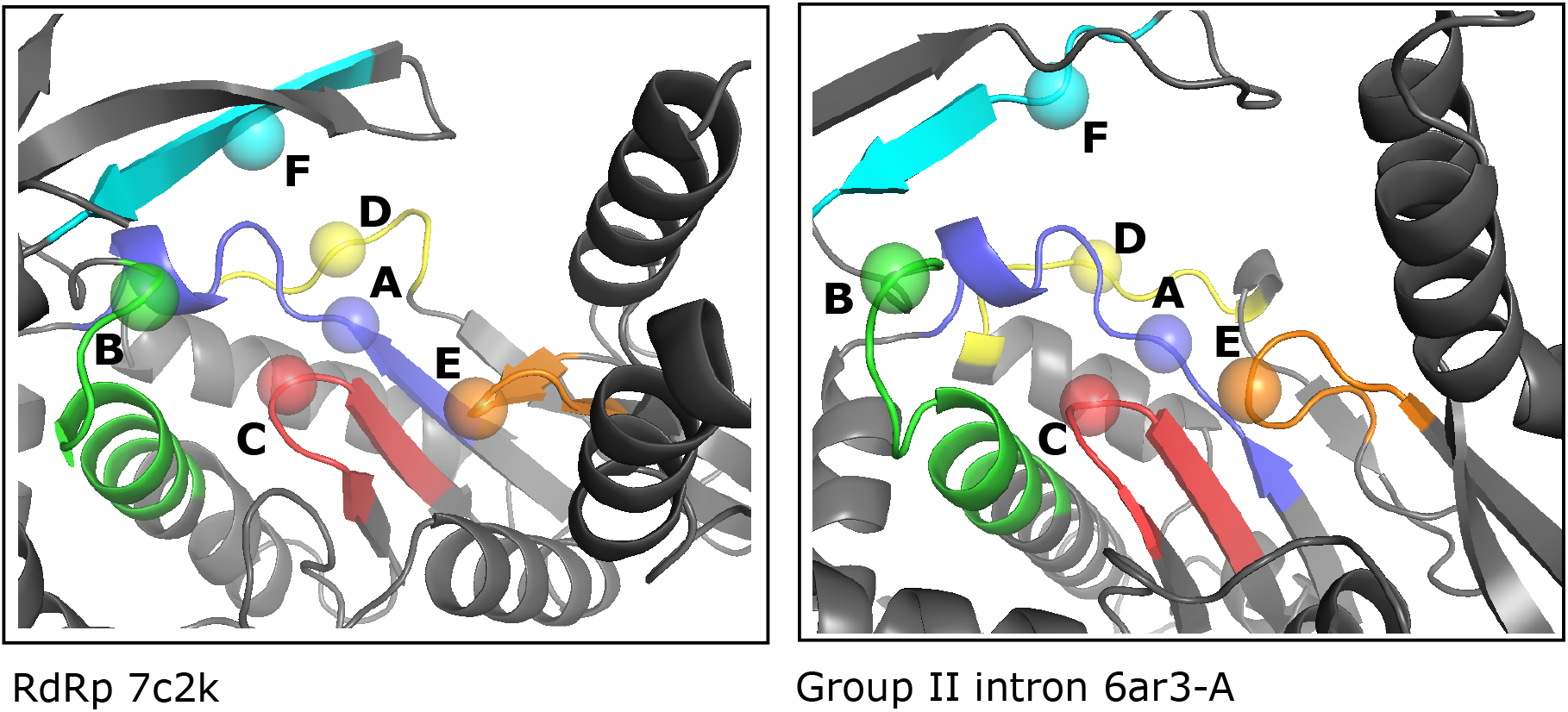
Structural alignment of viral RdRp to a cellular homolog. On the left is the palm domain of SARS-Cov-2 RdRp (PDB:72ck) aligned to *Geobacillus stearothermophilus* group II intron (right, PDB:6ar3) showing the essential conserved motifs A through F. Motif sequences are sometimes suggestive of assignment to viral RdRp or to a cellular homolog, but are never definitive.

**Table 4.**
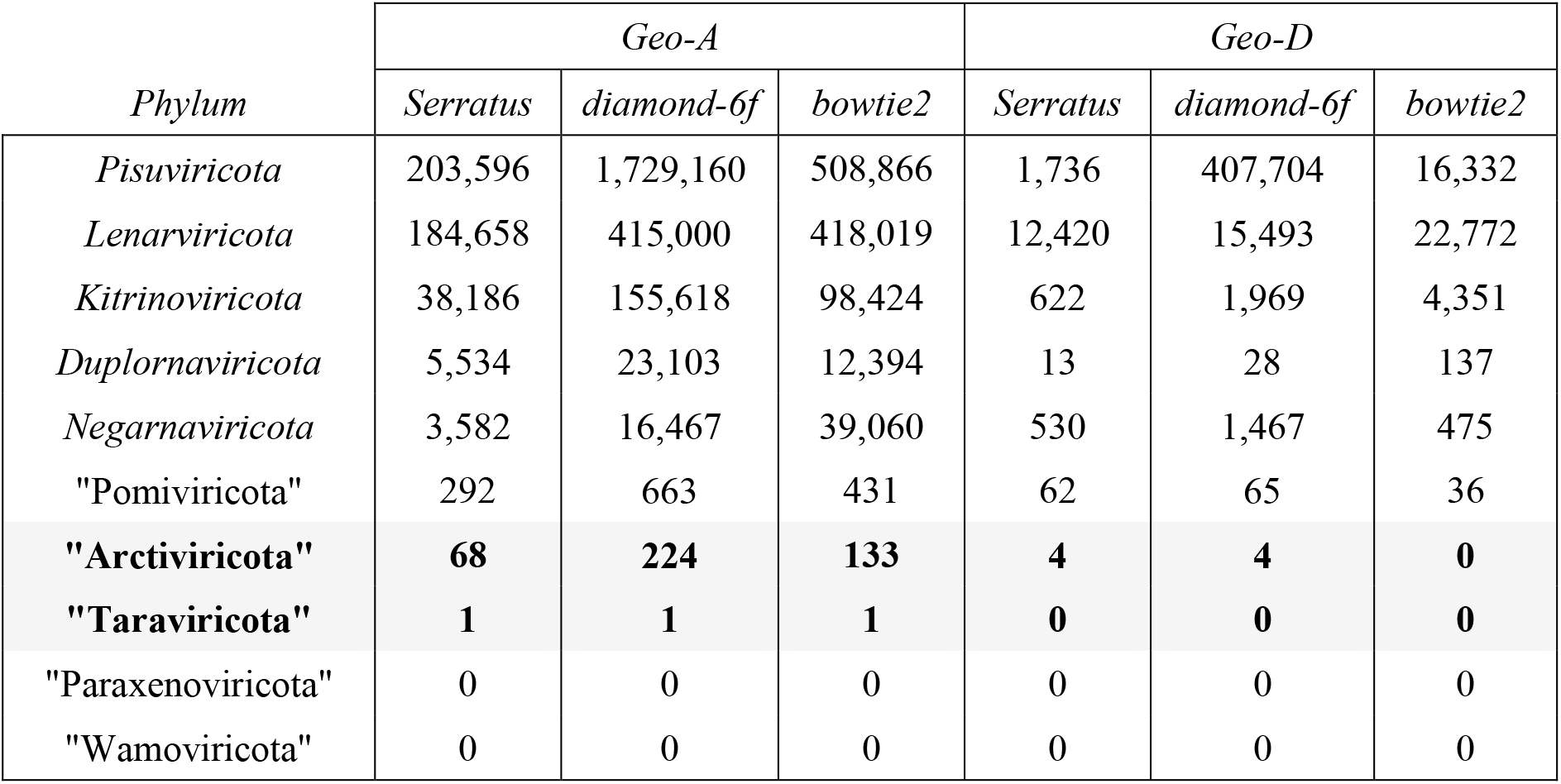
Numbers of reads mapped to geolocations A and D. This table shows the number of reads mapped per phylum using three different methods. Serratus is method (1), diamond-6f is method (3) and bowtie2 is method (4) as described in my main text under ‘Focused re-analysis of selected geolocations’. My results show exactly one read at Geo-A and zero reads at Geo-D mapped to “Taraviricota” according to all methods, contradicting Tara’s Fig. 4 which shows high abundance of “Taraviricota” at both locations. The apparent large discrepancy between different methods for *Pisuviricota* at Geo-D is due to cellular proteins in the full-length contigs which are trimmed in the mapping reference for Serratus; diamond is more sensitive to these alignments than bowtie2.

| *Phylum* | *Geo-A* |  |  | *Geo-D* |  |  |
| --- | --- | --- | --- | --- | --- | --- |
|  | *Serratus* | *diamond-6f* | *bowtie2* | *Serratus* | *diamond-6f* | *bowtie2* |
| *Pisuviricota* | 203,596 | 1,729,160 | 508,866 | 1,736 | 407,704 | 16,332 |
| *Lenarviricota* | 184,658 | 415,000 | 418,019 | 12,420 | 15,493 | 22,772 |
| *Kitrinoviricota* | 38,186 | 155,618 | 98,424 | 622 | 1,969 | 4,351 |
| *Duplornaviricota* | 5,534 | 23,103 | 12,394 | 13 | 28 | 137 |
| *Negarnaviricota* | 3,582 | 16,467 | 39,060 | 530 | 1,467 | 475 |
| "Pomiviricota" | 292 | 663 | 431 | 62 | 65 | 36 |
| **"Arctiviricota"** | **68** | **224** | **133** | **4** | **4** | **0** |
| **"Taraviricota"** | **1** | **1** | **1** | **0** | **0** | **0** |
| "Paraxenoviricota" | 0 | 0 | 0 | 0 | 0 | 0 |
| "Wamoviricota" | 0 | 0 | 0 | 0 | 0 | 0 |

### Abundance measure

Tara calculated abundances using CoverM, but the details are not completely specified. It is described as follows: “For the vertical coverage (i.e., for abundance estimation), reads that mapped at ≥ 90% ID over ≥ 75% of the read length were extracted using CoverM v0.2.0-alpha6, calculating the trimmed mean (tmean) for each contig. . . Only adjusted abundances of the ≥ 1-kb contigs were kept, and final abundances of the vOTUs were calculated by summing the adjusted abundances of the ≥ 1-kb contigs belonging to these vOTUs” (their Material and Methods under ‘Calculation of vOTU relative abundances’). No code is provided for calculating abundances from bowtie2 SAM files. Their supporting data does not include CoverM reports, or the calculated abundances per contig per sample or per vOTU per sample. It is not described how to calculate abundance of a “metataxon” or phylum from vOTU abundances, and a table of “megataxon” or phylum abundances per sample (needed to reproduce their Fig. 4) is also not provided.

Here, I measured abundance by the number of reads mapped because it is simple, transparent and adequate to identify “dominant” phyla. It is not stated whether Tara mapped reads from a given sequencing run to (a) the assembly for that run, or (b) to the combined set of contigs from all assemblies. Both alternatives are reasonable; (a) requires that a species is sufficiently abundant to be assembled before it is counted in a given sample, while (b) is more sensitive because it includes species which are rare in the reads for a given sample but are successfully assembled elsewhere. I could not do (a) because most contigs are not annotated with a sample or sequencing run, and therefore chose (b).

My result that ≪ 1% of reads map to “Taraviricota” and “Arctiviricota” at Geo-A and Geo-D necessarily implies that either Tara’s results or my results are unambiguously wrong, because the discrepancy between my figure and theirs is far too large to be explained by technical details of read processing or the choice of abundance metric. My results from read mapping over the full dataset correlate well with Tara’s own results for contig and vOTU frequencies, and I therefore believe that the balance of evidence suggests that Tara’s Fig. 4 is based on incorrectly calculated abundance values. This could be clarified if Tara annotates their contigs with sequencing runs, and makes their SAM files, CoverM reports and data analysis scripts available for independent review.

### Sequence and structure conservation in the RdRp domain

Viral RdRp belongs to the palm domain superfamily, with close homologs in several families of cellular palm domain proteins including group II introns. There are six essential motifs in the catalytic core of the polymerase palm domain (Lang et al., 2012), conventionally denoted by letters A through F which usually appear in order FABCDE in the primary sequence, though permuted variants are also known (e.g. (Gorbalenya et al., 2002)). A seventh motif G is sometimes included; I do not consider it further here because its structural conservation is less clear. Motifs F, A, B and C each have one conserved catalytic residue (F=ARG, A=ASP, B=GLY and C=ASP). Within a phylum, the A, B and C motifs are sufficiently well conserved to be recognizable by sequence (Babaian and Edgar, 2021), while the remaining motifs are more variable in primary sequence. Between different phyla, motifs are generally not recognizable by primary sequence similarity with the exception of GLY-ASP-ASP in motif C which has a few common variants such as SER-ASP-ASP in SARS-Cov-2. All six motifs are found in all solved structures for palm domain polymerases and reverse transcriptases (see (Lang et al., 2012) for motif coordinates in structures known in 2012). They are well conserved in structure, aligning well between viral RdRp and non-viral homologs including group II introns, as shown in my Fig. 4. If six new virus phyla can be discovered in Tara’s data, then surely one or more new families of group II introns, or some other close cellular homolog, could also be discovered. Given that the diversity of palm domain proteins is known only sparsely at the present time, it is simply not possible to distinguish with certainty a highly diverged RdRp from, say, a highly diverged group II intron by sequence and structure alone.

### Distinguishing viral RdRp from close homologs

In their Materials and Methods under ‘Evaluation of authenticity and completeness of putative virus RdRps’, Tara describe their protocol as follows: “Hits longer than 100 amino acids with a best match to an RdRp HMM and with a bitscore ≥ 30 were kept as true positives for proteins containing the virus RdRp domain. Lower-scoring hits were manually inspected for presence of the seven canonical RdRp domain motifs. In total, 44,779 contigs encoding putative virus RdRps were detected by these identification and curation processes.” While this procedure seems reasonable, it is not definitive—certainly not to the standard that should be required for identifying novel phyla—and some residual false positive rate should be expected. “Best match to an RdRp HMM” is a top-hit / nearest-neighbor test which may give false-positive assignments because top hits are not necessarily evolutionary neighbors (Koski and Golding, 2001). Using a bit score threshold is unconventional, why were E-values not used? A bit score of 30 is low and typically corresponds to a high E-value; the corresponding E-value should therefore have been quoted and the choice of bit score threshold should have been explained. Presumably, some or all of the claimed novel phyla are among the most diverged and hence lowest-scoring hits, but absent documentation it is not possible to verify this. “Manual inspection” cannot reliably identify catalytic motifs in highly diverged proteins because the motifs are conserved in structure but not in sequence. Even within a known phylum, in my experience it is often difficult or impossible to identify D, E and F motifs without structure. While rules of thumb can be applied to motif sequences, e.g. GLY-ASP-ASP in motif C is characteristic of RdRp while ALA-ASP-ASP is characteristic of reverse transcriptases, there is to the best of my knowledge no reliable method for distinguishing viral RdRp from a cellular homolog given sequence and structure alone because there are exceptions. For example, AOY33888 (RdRp of squash vein yellowing virus) has ALA-ASP-ASP in its C motif, and conversely WP 014123481 (group II intron of bacterium *Tetragenococcus halophilus*) has GLY-ASP-ASP. Tara should therefore have documented the criteria they used to decide whether motifs were indicative of RdRp, and provided a list of low-scoring hits together with their inferred motifs and functional assignments, to enable independent review.

### Cellular proteins identified in Tara’s putative viral contigs

I aligned Tara’s RdRp+ contigs to the NCBI non-redundant protein database using diamond, collecting the top hits only. 510 contigs had hits to non-viral proteins with *E* < 10 ^−9^ (My Supplementary Table S1), 137 of which had aa sequence identity > 90%. Many of these proteins are associated with cellular marine organisms including algae, plankton and dinoflagellates. If an RdRp-like sequence is found in a contig with an identical or highly similar match to a cellular protein, this favors the hypothesis that the contig was derived from a cellular transcript rather than a viral genome or transcript. The RdRp-like sequence can then tentatively be identified as a cellular RdRp homolog such as a group II intron. Other explanations are possible, including an incorrectly assembled contig comprising a viral/cellular chimera, a recent virus insertion into a host, or a recent capture of a host gene by a virus, but here the parsimonious explanation is a correct non-viral contig containing a cellular RdRp homolog, or at a minimum this should be the null hypothesis before claiming a novel virus, and the burden of proof should therefore be on Tara to rule out other explanations. Further, if the full-length contig is used for read mapping and virus abundance estimation, the cellular protein may cause spurious inflation of abundance due to mapping of non-viral reads onto the contig. An example is shown in my Fig. 5, which annotates a contig which Tara assigns to “Taraviricota” and meets their criteria for a reference sequence, i.e. it is > 1knt and has an estimated RdRp domain completeness of 100% according to their supplementary tables. This contig contains a 59% identity match to a dinoflagellate protein with E = 10^-31^.

**Figure 5.**
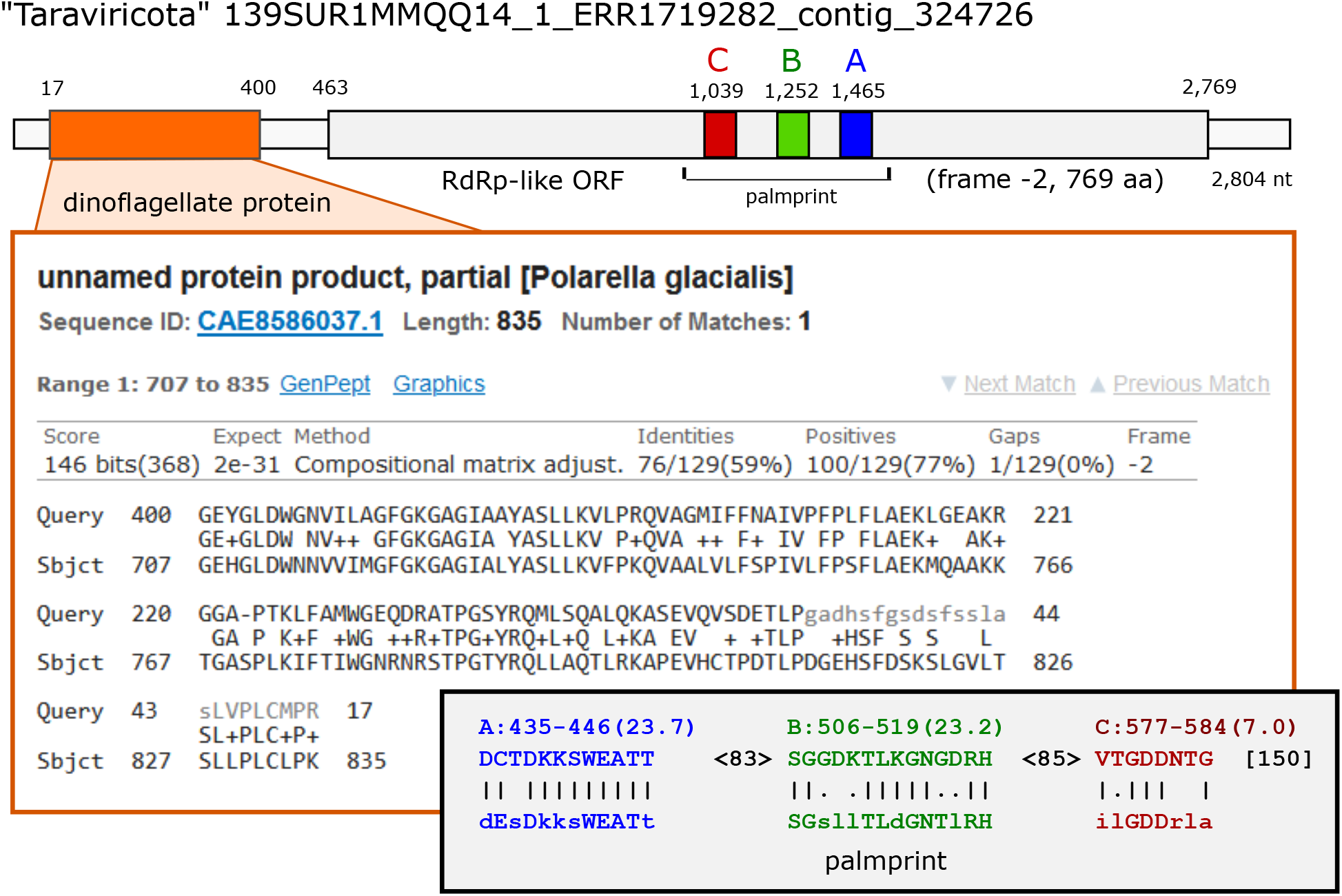
“Taraviricota” contig with dinoflagellate protein. This contig is 2,804nt. It is assigned to “Taraviricota” with 100% estimated RdRp domain completeness by Tara Table S6. Positions 7 through 400 are shown as aligned to *Polarella glacialis* protein CAE8586037.1 by BLASTX. Positions 463 through 2,769 are an RdRp-like ORF on the negative strand. Predicted motifs A, B and C according to palmscan are shown in the lower inset. Numbers shown at top are nt coordinates counting from 1 at the beginning of the contig. Predicted motif coordinates by palmscan are aa positions counting from 1 at the start of the ORF. More than 500 of Tara’s putative viral contigs have top hits to cellular proteins with *E* < 10^-9^, many of which have 100% aa identity (my Supplementary Table S1).

### Predicted RdRp structures for Tara’s “phyla”

According to Tara’s Material and Methods under 3D structure network analysis, they “predicted the 3D structures for the new megataxa from their representative primary amino acid sequences (the longest sequence with no ambiguous residues (i.e., no ‘X’s in the primary sequence) per megatxon) using Phyre2 in the ‘Normal’ mode”. They deposited five structures, one for each “phylum” in cyverse: https://de.cyverse.org/anon-files//iplant/home/shared/iVirus/ZayedWainainaDominguez-Huerta_RNAevolution_Dec2021/Predicted_3D_Structures/.

I downloaded the pdb files for these structures and aligned them to SARS-CoV-2 RdRp (PDB:7c2k) using pymol (https://www.pymol.org). The predicted structure for “Taraviricota” aligned well and in my judgment appears consistent with a polymerase or reverse transcriptase in the palm domain superfamily. However, structures for the other four “phyla” are obviously truncated and malformed. My Fig. 6 summarizes their alignments to CoV. These four predicted structures range in length from 142 aa (“Arctiviricota”) to 211 aa (“Wamoviricota”) and are thus much shorter than a complete RdRp domain which ranges from a minimum of more than 400 aa to a maximum of > 1,000 aa (Lang et al., 2012). The catalytic cores of the predicted palm domains are truncated such that one or more essential motifs are missing, most obviously in “Pomiviricota” which lacks motifs A, B and C and F. In “Wamoviricota” and “Parexenoviricota” roughly half the catalytic core is present but obviously malformed, as shown in my Fig. 7. For example, in “Parexenoviricota”, one strand of the anti-parallel beta sheet of motif C is replaced by one helix turn and in “Wamoviricota”, the alpha helix of motif B, which has five or more complete turns in all known structures, has a single turn followed by a loop. Therefore, if these predicted structures are substantially correct, they are sufficiently different from known palm domain polymerases to contradict the hypothesis that they are viral RdRp. Conversely, if the predictions have substantial errors and the true structures closely resemble known viral RdRps, then the predicted structures are sufficiently defective to undermine inferences of function and phylogenetic relationships by comparison with solved palm domain structures.

**Figure 6.**
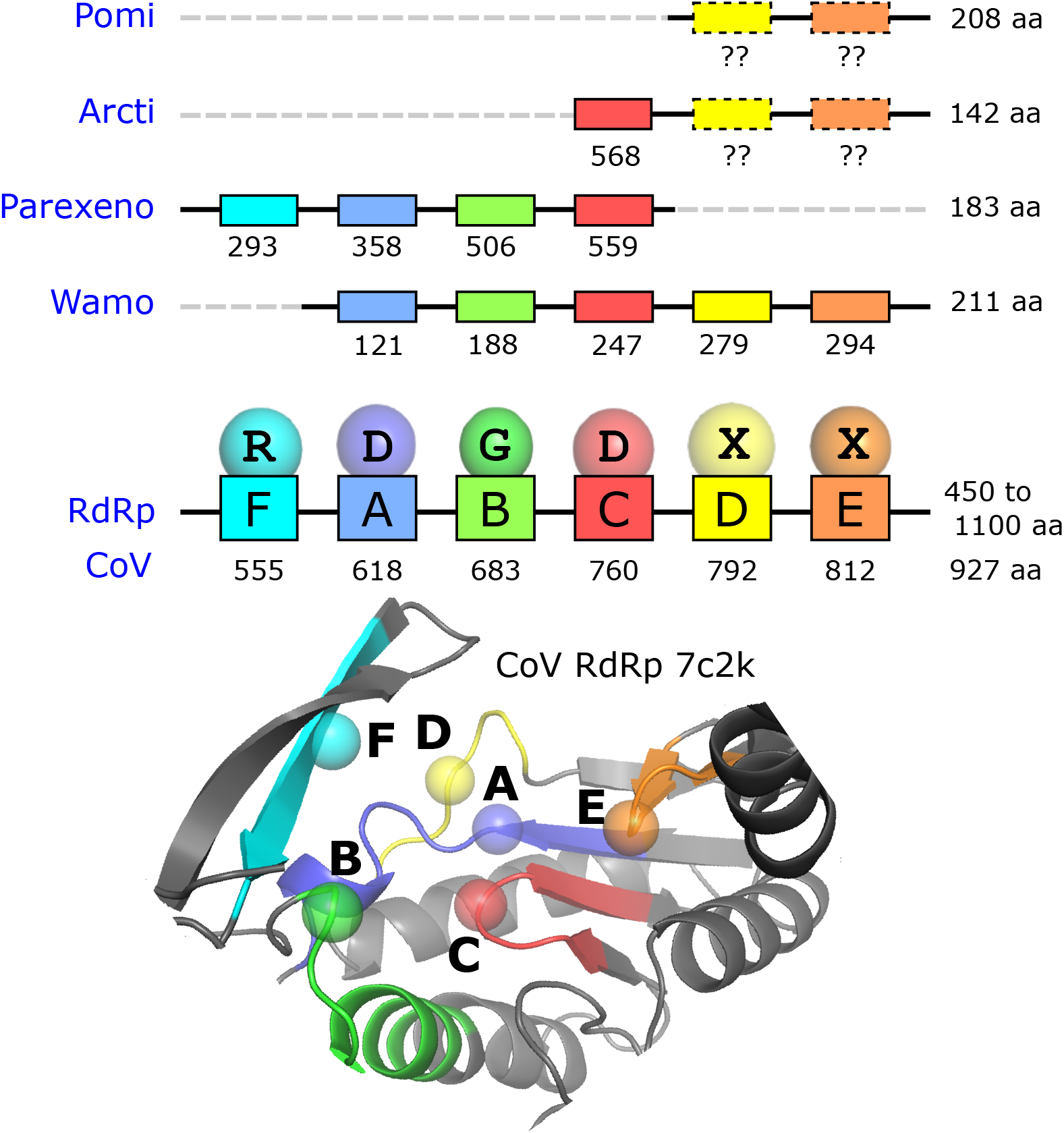
Structural alignments of Tara predicted structures to solved viral RdRp. Four of Tara’s predicted structures are aligned to SARS-CoV-2 RdRp (PDB:72ck). The CoV structure cartoon is colored to show the six essential motifs FABCDE, with catalytic residues indicated by spheres. Above the cartoon is a schematic showing inferred motif positions in each structure as residue numbers in the pdb files (note that residue numbers in the Tara pdb files are not 1-based). Chain lengths are given at the right-hand side. Phylum names are abbreviated to Pomi=“Pomiviricota”, Arcti=“Arctiviricota” etc. All Tara structures except “Taraviricota” (not shown) are truncated such that one or more motifs are missing. Motifs D and E in solved structures did not align well to the corresponding regions in Pomi and Arcti and I was therefore not able to verify the presence of these motifs (see my Fig. 7).

**Figure 7.**
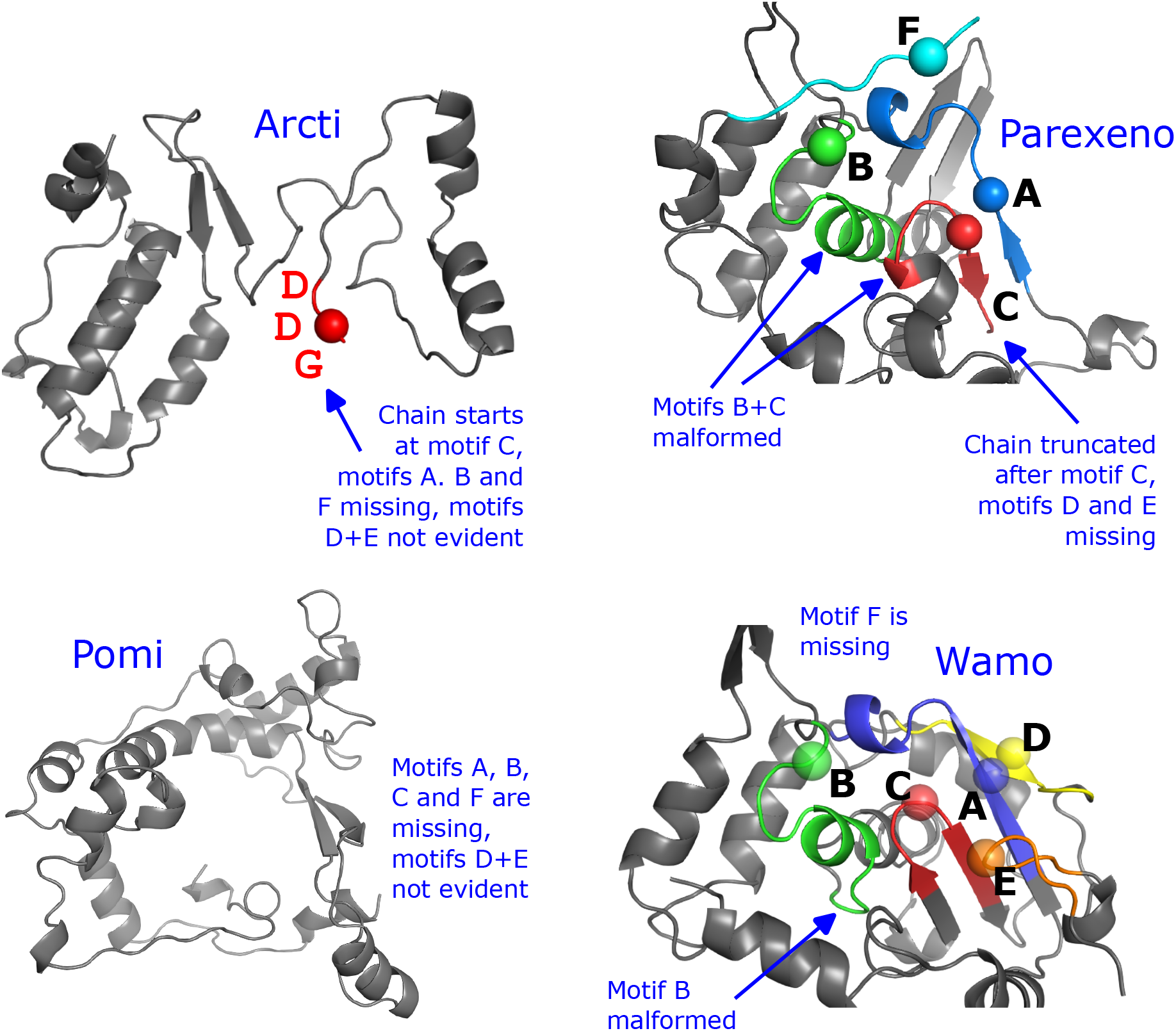
Truncated and malformed palm domains in predicted structures. The figure shows the complete structures of Arcti, Parexeno, Pomi and Wamo as archived in Tara’s cyverse repository (phylum names without “viricota” for brevity). None of these structures has a complete palm domain. The Pomi structure is truncated such that the entire chain aligns approximately to a region starting after motif E, which implies that all conserved motifs are deleted if in fact they are present. In Arcti, the first three residues of the chain are GLY-ASP-ASP which align approximately to motif C, but the characteristic conformations of motifs C, D and E are not evident. Parexeno and Wamo align to roughly half of the palm domain, but the catalytic core is obviously malformed (my Fig. 8).

### Annotation of predicted structures in Tara’s Fig. S5

My analysis of the predicted structures conflicts with Tara’s Fig. S5, reproduced with added notes as my Fig. 9. Tara marks 12 motifs as present which in fact are unambiguously absent due to truncation of the palm domain. Motif E is annotated as “naturally absent” in *Lenarviricota* PDB:3mmp-G, but in fact this essential motif is found at residue 397 as shown in my Fig. 10.

**Figure 8.**
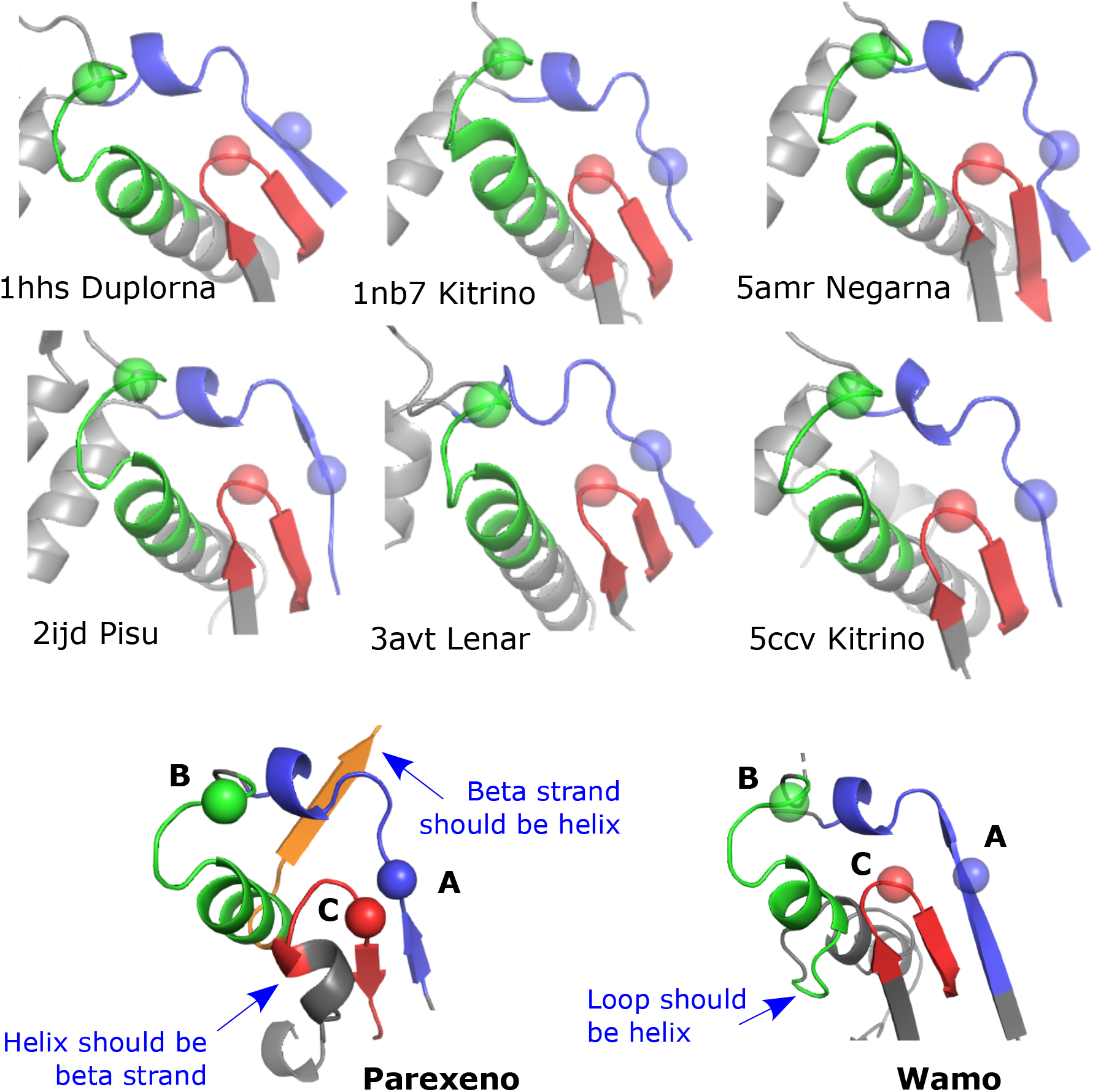
Malformed catalytic cores of “Parexenoviricota” and “Wamoviricota” structures. The figure shows the “palmprint”, i.e. the palm domain segment from motif A to motif C (Babaian and Edgar, 2021), in Parexeno and Wamo with six representative solved structures for comparison (phylum names without “viricota” for brevity). The alpha helix of motif B should have five complete turns, but in Parexeno there are three and in Wamo only one. In Parexeno, the three helix turns are followed by a beta sheet which is perpendicular to three helix turns at the corresponding primary sequence positions when aligned to solved structures. In Wamo, the single turn is followed by a loop. In Parexeno, the characteristic antiparallel beta sheet of motif C is replaced by a single-turn helix adjacent to a beta strand.

**Figure 9.**
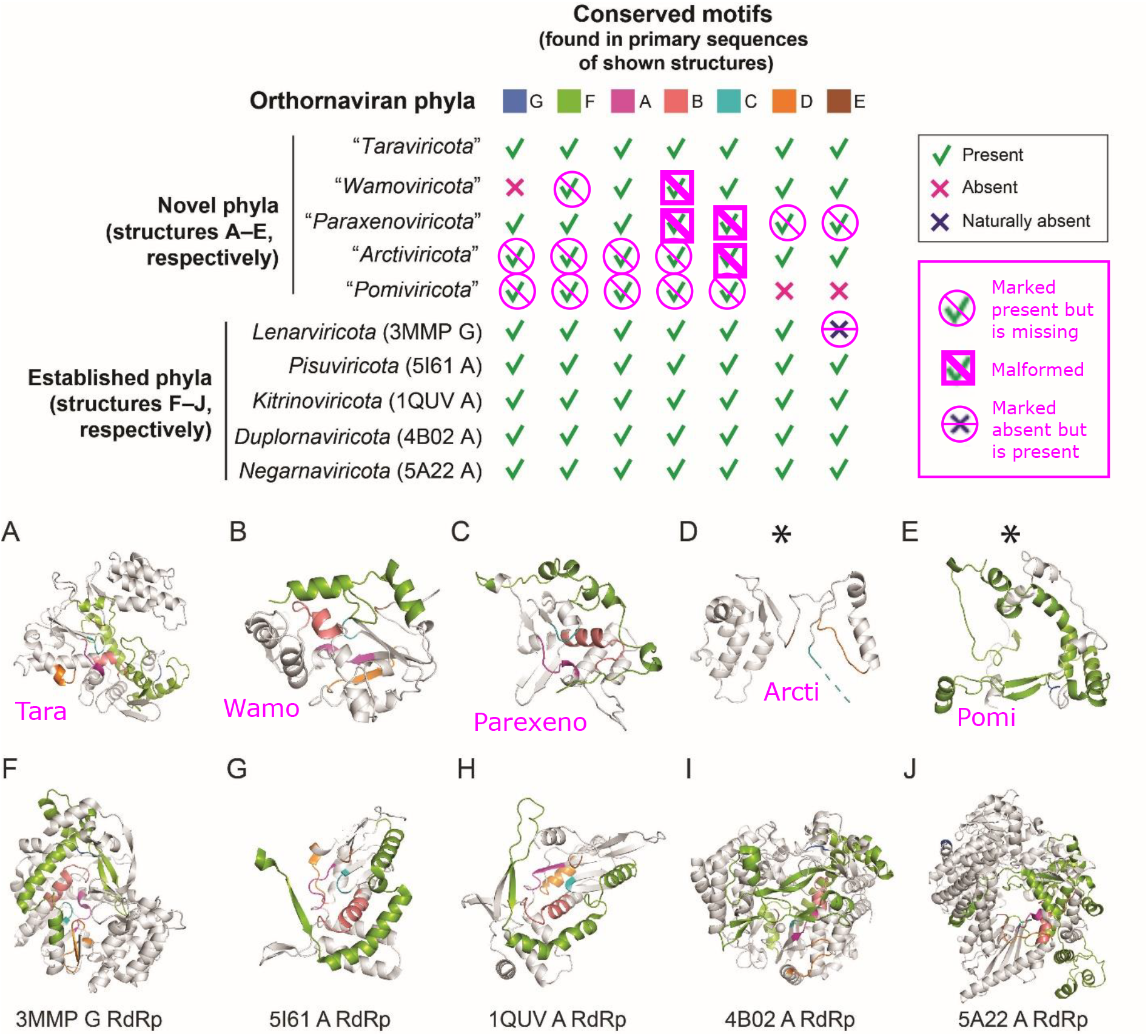
Conflicting motif identification in Tara structures. The figure shows Tara’s supplementary Fig. S5, with my notes added (magenta color). I find 12 motifs marked as present by Tara to be definitively absent due to truncation of the domains. Four motifs are present but obviously malformed. Essential motif E is annotated as “naturally absent” in PDB:3mmp by Tara, but in fact this motif is present at residue 397 as shown in my Fig. 10.

**Figure 10.**
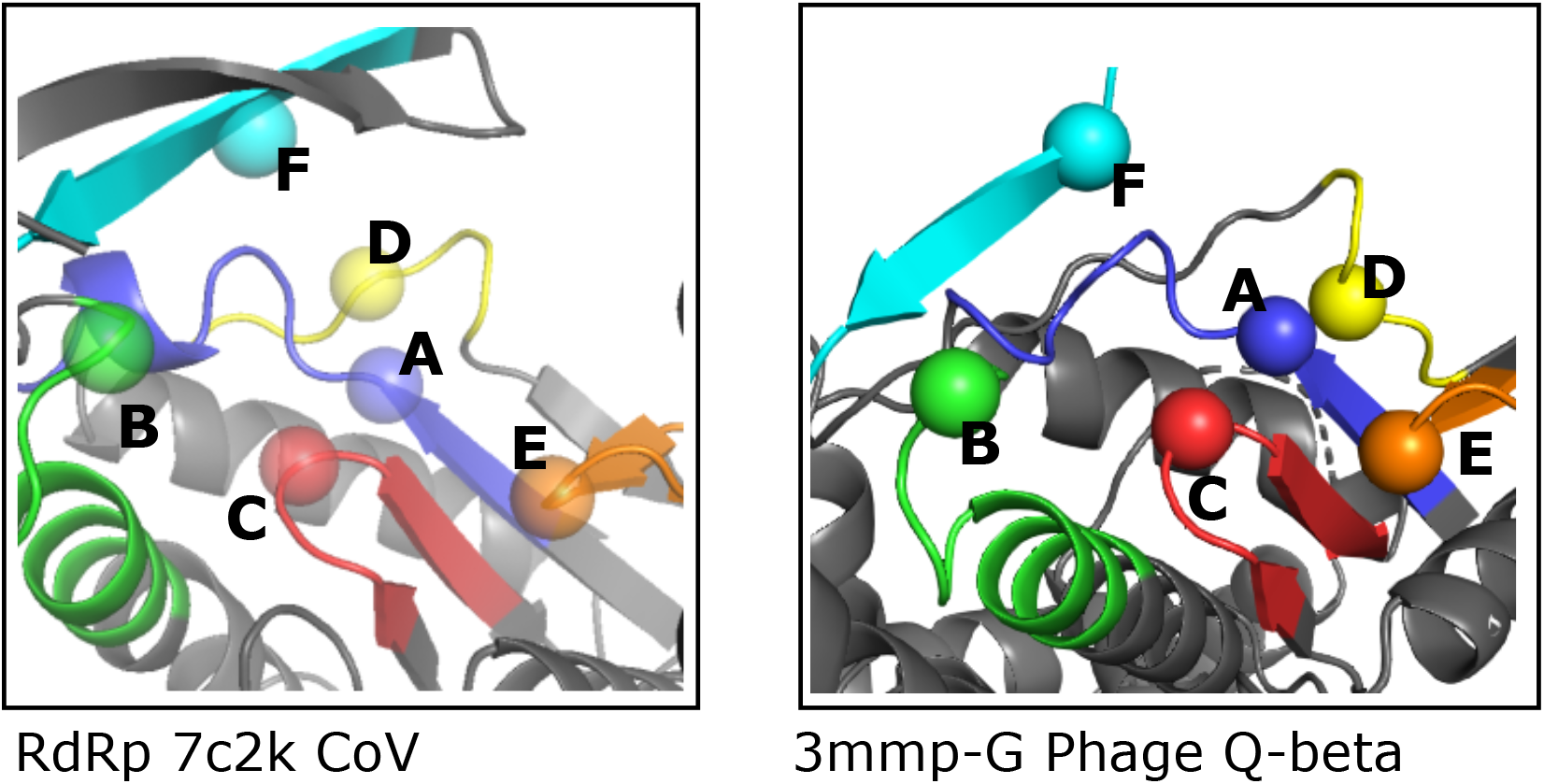
Conserved motifs in phage Qβ. The figure shows the palm domain of phage Qβ (PDB:3mmp-G, right panel) aligned to SARS-CoV-2 RdRp (PDB:72ck, left panel). Motif E is located at residue 397 in 3mmp-G, contradicting the annotation in Tara’s Fig. S5 that this motif is “naturally absent”.

### Tara’s structure network

Tara’s Fig. 3B shows a “structure network” which, according to the figure caption, is informative for “…inferring the early history of orthornavirans”. As described in their Material and Methods under “3D structure network analysis”, pair-wise structural alignments were constructed using Matras v1.2 (Kawabata, 2003). For each pair, the superfamily similarity reliability score was used as a distance. The network was generated by cytoscape (Shannon et al., 2003) from these distances, using the using the “Edge-weighted Spring Embedded” layout. No citations to previous applications of networks to phylogenetics are given, no justification is given from theoretical considerations, and no validation is reported which assesses the accuracy of inferences made from this type of network. The remarkable claim that this *ad hoc* method is informative for deep phylogenetic inference is thus not supported by any evidence. Further, the procedures and criteria used to make inferences from the network are not explained; conclusions are simply presented as *faits accomplis*.

Why Matras scores and cytoscape? Could DALI (Holm and Sander, 1995) Z-scores or TM-align (Zhang and Skolnick, 2005) TM scores be used instead? Could the “Compound Spring” or “Prefuse Force” layouts be used, or is “Edge-weighted Spring” superior for some reason? Could the network be generated using Gephi (gephi.org) or Wandora (wandora.org)? More generally, which combinations of similarity scores and network construction methods are valid, and which are invalid? What objective criteria enable phylogenetic inferences from these networks? What types of hypotheses can be robustly confirmed or contradicted? How can the accuracy of these conclusions be assessed? In the absence of a framework for delivering evidence-based answers to questions like these, this type of method cannot support a rational approach to classification.

### Tara’s “megataxonomy”

Tara’s classification method is sketched in my Fig. 11. Translated RdRp sequences from Tara contigs were combined with RdRps from Yangshan Deep-Water Harbour and GenBank. The combined RdRps were reduced to 13,109 non-redundant centroid sequences at 50% identity by UCLUST (Edgar, 2010). The centroids were then clustered by MCL (Enright et al., 2002) from a matrix of pair-wise BLASTP bit scores, giving 19 “megaclusters”. Each megacluster was assigned a “megataxon” based on the majority taxon or taxa according to annotations of its GenBank sequences.

**Figure 11.**
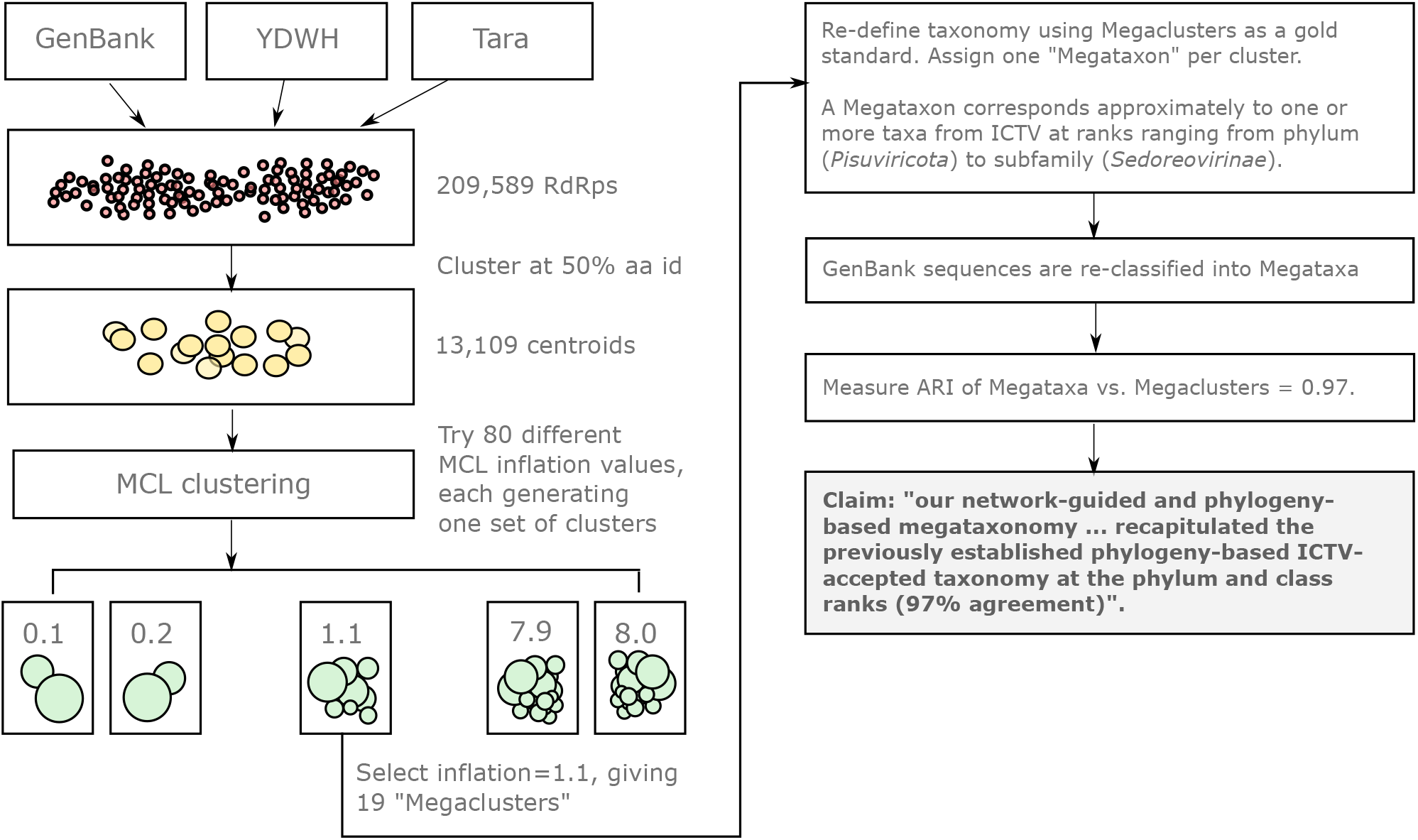
Tara’s “megacluster” workflow and incorrect claim of 97% agreement with ICTV taxonomy. RdRp amino acid sequences were collected from GenBank, Yangshan Deep-Water Harbour (YDWH) and Tara contigs. These were clustered twice: first at 50% aa identity, giving 13,109 centroids. Centroids were clustered by MCL using BLASTP bit scores as a similarity measure, giving 19 “megaclusters” at inflation=1.1. Motivations for design choices including MCL for clustering, BLASTP for similarity, and inflation value of 1.1, are not explained. For each “megacluster”, the consensus taxon or taxa of its GenBank sequences (the “megataxon” of the cluster) was identified. If the justification is entirely post-hoc, as is apparently the case, this procedure should be expected to over-fit because all labeled data is used in training with no hold-out for validation. “Megataxon” ranks range from phylum to sub-family (my Table 1), and with three exceptions (*Chrymotiviricetes, Vidaverviricetes* and *Allassoviricetes*), “megaclusters” do not attempt to classify at ICTV class rank. Tara measured agreement by adjusted Rand index (ARI) between “megaclusters” and “megataxa”, not percentage agreement between megataxa against ICTV phylum and class as stated in their claim.

In taxonomy generally, and per ICTV rule 3.3.1 specifically, monophyly is a fundamental requirement for defining taxa. However, it is textbook knowledge that clustering cannot reliably infer phylogenetic trees or monophyletic groups, with one exception based on unrealistic assumptions (the correct tree can be reconstructed by UPGMA if the molecular clock is ultrametric and true evolutionary distances can be calculated). This result was established in the 1960s in the first literature to consider mathematical and algorithmic applications to phylogenetics, and has been universally accepted since then. See Chapter 10 in ‘‘Inferring Phylogenies” (Felsenstein, 2004) for history, methodological survey and references.

MCL is a generic clustering method, and therefore cannot reliably predict monophyletic groups. Adopting an approach of this type for classification would require abandoning monophyly as a standard, without providing alternative classification principles. If a future study finds that MCL with inflation 1.1 based on BLASTP bit scores produces quite different clusters from Tara’s when new data is added, which seems inevitable, then how to proceed? Should Tara’s “megataxa” be abandoned? Should a different clustering method or similarity score be tried? Tara offers no guiding principles for answering such questions.

### Phylogenetic tree

Tara’s Fig. 3A shows a phylogenetic tree obtained by MSA and ML. Inspection of Tara’s phylum-level MSA and tree (Global_RdRp_Tree_sequence.fasta.aln and Global_RdRp_Tree_newick.txt in Tara’s DRYAD data repository) show that exactly one sequence was included for four of the five established ICTV phyla. It is therefore impossible by construction for any other sequence to place within the subtree of an ICTV phylum, with the exception of *Duplornaviricota* which has nine sequences. Thus, this tree cannot support a hypothesis that novel sequences fall outside known phyla.

### Claim of 97% accuracy for taxonomy assignment

Tara claimed that their MCL clusters “nearly completely recapitulated the previously established phylogeny-based ICTV-accepted taxonomy at the phylum and class ranks (97% agreement)” on the basis of the ARI=0.97 value reported in their Fig. 1B. However, the agreement underlying this claim is not the fraction of correct classifications of phylum and class with GenBank taxonomy annotations as implied by the statement; rather, it is the adjusted Rand index obtained by comparing “megataxa” and megaclusters. “Megataxa” do not correspond to phylum and class separately or to phylum and class together; rather they are a heterogeneous collection of phyla, sub-phyla, polyphyletic groups (e.g. “Lenarviricota, others”), classes, unassigned ranks (e.g. “Wei-like”), and sub-families. With three exceptions (*Chrymotiviricetes, Vidaverviricetes* and *Allassoviricetes*) “megataxa” cannot be used to classify to ICTV class rank.

This *ad hoc* classifier was tuned to all available training data, which surely results in extreme overfitting because, as with the 3D structure network, there are no stated prior restraints on variations which can be tried, including the choice of distance metric, choice of clustering method, choice of clustering parameters (e.g. inflation value), and choice of ICTV taxa assigned to a cluster. The number of possible variations is thus astronomical, and the opportunities for over-fitting are therefore for all practical purpose unlimited. Given that a high degree of over-fitting should be assumed, it follows that the agreement between megaclusters and “megataxa” is not predictive of agreement when classifying new, unlabeled sequences (Santos et al., 2018), even if it is accepted that “megataxa” defined by this approach should supplant ICTV taxa.

A more realistic and informative assessment of classification accuracy would be obtained by randomized hold-out validation using ICTV taxa as the standard of truth as opposed to “megataxa”.

## Conclusion

The Tara authors’ own summary data show that their claimed novel “phyla” comprise a small fraction of vOTUs and a small fraction of their contigs. My results show that the evidence for viral origin of their putative novel RdRps is equivocal, and regardless the relevant putative RdRp-containing contigs account for only a small fraction of the reads. Together, these results comprehensively contradict Tara’s claim that new phyla are “dominant in the oceans”. In fact, known phyla account for a large majority of their contigs, vOTUs and reads, and known phyla thereby dominate the Tara Oceans RNA virome.

### Code and data availability

Code and data are deposited at https://github.com/rccdgar/tara_oceans and https://zenodo.org/record/7194888.

